# From Flux to Function: Extracting Mechanistic Insights from Ion Channels via I-V and I-μ Analyses

**DOI:** 10.64898/2025.12.01.691660

**Authors:** Hannah Weckel-Dahman, Ryan Carlsen, Alexander D. Daum, Jessica M.J. Swanson

**Affiliations:** Department of Chemistry, University of Utah, Salt Lake City, UT, 84112 – United States of America

**Keywords:** Multiscale Responsive Kinetic Modeling (MsRKM), Ion Channels, Electrophysiology, Current-Voltage (I-V), Electrochemical Gradients, Ohmic Flux, Rectification

## Abstract

The molecular origins of ion channel current–voltage (I–V) relationships are often unclear, obscured by ensemble averaging in experimental analysis and persistent underestimation of single-channel ion currents in simulations. Here, we present a mathematical framework that relates ion channel properties to experimentally measured I–V and current–concentration (I–μ) relationships. By accounting for how rates change in response to electrochemical conditions in a multistate kinetic model of systems with sequential binding sites, this approach demonstrates how the spatial arrangement of sites and transition states, together with rate asymmetries in ion uptake, transfer, and release, manifest in distinct open-channel current and conductance profiles. Varying these properties in model systems reveals a molecular basis for understanding rectification and non-ohmic open-channel flux. Application to more realistic models fit to I–V curves for the Shaker K_v_ channel demonstrates that these mechanistic trends hold in heterogeneous systems, suggesting a (potentially) transferable paradigm for open channel flux in channels and transporters with two or more sequential binding sites. Together, these results establish a theoretical framework for open channel current and foundation for mechanistically interpreting experimental I–V and I–μ assays.

## 1. INTRODUCTION

Ion channels allow the dissipation of electrochemical gradients through selective permeation of ions across membranes, generating neuronal and cardiac electrical activity, transducing chemical signals, and maintaining osmotic and volume homeostasis.^1-4^ Two central areas of ion channel research aim to understand 1) how channels gate—open and close in response to voltage, ligands, mechanical force, or metabolites—and 2) how they support rapid, selective ion permeation once open. This work explores the mechanistic underpinnings of the latter—open channel current. Although gating was initially more difficult to study, notable advances over the past several decades have significantly advanced our understanding gating mechanisms, continuing to support progress toward pharmacological correction of abnormal ion channel function.^5-7^ Concurrently, progress has been made toward understanding open-channel conduction by integrating simulation-based insights with electrophysiological data, often in the form of I-V curves. Despite these advances, unraveling the molecular mechanism of open-channel current remains a central challenge—largely due to inconsistencies between simulations and experiments. Gaps in our understanding of open-channel current warrant continued attention, not only from a fundamental perspective but also because certain disease states are definitively tied to mutants that vary only in open-channel current.^8-10^

Historically, efforts to understand the molecular determinants of open-channel currents have benefitted from the complementary strengths of experimental measurements and computational analysis. Early molecular dynamics (MD) simulations provided valuable atomic-level insights into open-channel ion conduction, ion selectivity, hydration, and conduction pathways.^11-13^ One of the remaining frontiers in ion channel research is reconciling experimentally-observed conduction rates with those predicted by MD simulations—a challenge that has driven methodological innovation over the past two decades. Recent advances, including the integration of polarization effects^14^ and simulation post-processing via Markov models,^15^ represent important milestones in this progression. Despite these developments, consistently matching experimental I-V data remains an outstanding challenge. Additionally, extracting the underlying transport mechanisms from these methods requires flux pathway simplification that may inadvertently modify channel response.^16^

An alternative approach to explain open-channel ion current is kinetic modeling,^17-21^ which translates the ensemble, macroscopic property of ion flux (provided by I-V curves) into mechanistic insight by representing ion transport through a protein as a network of discrete metastable states connected by transition rates. Solving the kinetic master equation resolves flux through this network for all nonequilibrium conditions for which the included transition rates apply, opening the door to understanding ion channels under physiologically-relevant nonequilibrium conditions.^16,22,23^ To accomplish this, kinetic models must account for how rates change in response to those conditions. Electrical gradients are expected to alter rates via the underlying ion transport free-energy landscapes in at least two ways: via condition-induced conformational changes, including those that enable gating, and via voltage-induced alteration of charge stability across the membrane.^13,24-27^ Although the former can take many forms, the conformational changes associated with gating can be included in kinetic models, linking open and closed channel states.^20,28-30^ For the latter, voltage influences the stability of charged species along the channel pore, changing not only the relative stabilities of ion binding sites, but also the rates of transitioning between them. Recent work has demonstrated how this can in turn change the relative contributions of different transport pathways.^16,23,31,32^ Capturing these voltage-dependent effects provides a foundation for extracting the ion transport mechanisms that result in I-V curves.

From the name I-V curve, Ohm’s law is immediately considered with the implication of linear resistance. However, when looking at experimental results, I-V curves have a variety of curvatures, both superohmic and subohmic.^33-35^ Research on nanopores has indicated that superohmic curvature stems from enthalpic barriers, while subohmic curvature indicated entropic barriers.^36^ In ion channels, however, such a simple distinction is unlikely to apply since transport barriers result from a complex interplay of molecular interactions, rather than the primarily geometric or electrostatic constraints that govern diffusional dynamics in nanopores. Closely tied to variable I-V curvature is the observation that many ion channels exhibit a preferential direction of conduction (rectification), even in the fully open state. This rectifying behavior can be attributed to blocking ions or asymmetric charge distributions in some cases,^34,37,38^ but often has an origin that is not well understood. Lastly, I-V curvature can be influenced by the interplay between voltage-driven flux and ion concentrations, the balance of which depends heavily on the mechanism^33^ in a manner unique to each channel.

In this study, we explore how the structural features of ion channels lead to different flux profiles in open-channel conductance. Using responsive kinetic models that incorporate electrochemically dependent rates, we map the effects of channel properties on flux under a range of electrochemical conditions. In addition to I-V comparisons, we further demonstrate the rich physics that is uniquely captured in flux driven only by concentration gradients, I-μ curves, providing relationships that are again reflective of channel properties. We find that the locations of ion binding sites, transitions states, and relative rates of ion uptake, transfer, and release manifest in distinct conductance and current profiles. In turn, distinct conductance and current profiles reveal the location of the flux limiting step and explain the origin of rectification and non-ohmic flux. We then verify the extension of these findings to more realistic channel models with kinetic solutions that were previously fit to voltage-driven current profiles for the Shaker K_v_ channel. Collectively, these findings establish a paradigm for understanding the physical origins of I-V and I-μ relationships.

## 2. METHODS

### 2.1 Biomolecular Ion flux as a Function of Electrical and Chemical Potentials

The transport of charged species across a biological membrane separating an electrochemical gradient is driven by the magnitudes of the voltage and chemical gradient. Although the voltage decreases across the membrane in a roughly linear manner, the actual voltage drop is specific to the transport pathway and hence, for ion channels, to the protein environment. The fraction of the membrane potential experience by ions moving along the transport pathway—typically a reaction coordinate along the membrane normal (z)—can be quantified using a dimensionless coupling factor (*φ*(*z*)).^26,27,39,40^ This dimensionless coupling factor shifts the transport free energy profile, and hence transition activation energies, for the range of transmembrane potentials that do not induce a significant conformational change (e.g., voltage-induced gating), as derived in SI Section S1.1. This leads to an expression for the net flux of ions through a system^23^:

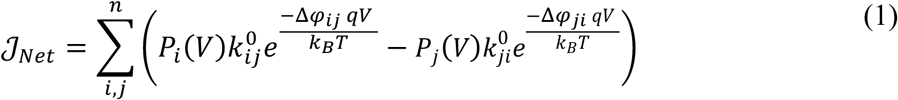

where i and j are linked pairs of states such that every pair up to the nth pair share a common transition state, either separating ion uptake and release or separating consecutive binding sites; k_ij_^0^ is the rate of an ion moving from state i to state j with no applied voltage; P_i_ and P_j_ represent the populations of states i and j, respectively; Δ*φ_ij_* = *φ_ij_* (*z*^‡*i,j*^) − *φ_ij_* (*z*^*bs_i_*^) is the difference between the fraction of the membrane potential at a transition state (*z*^‡*i,j*^) and stable binding site (*z*^*bs_i_*^) of site i (or bulk solution for uptake); q is the charge of the ion (or charged substrate); k_B_ is the Boltzmann factor; and T is temperature. Note that Δ*φ* effectively measures the voltage sensitivity for a given transition. Bulk intracellular and extracellular ion concentrations ([ion]_intra | extra_) are then folded into the net flux by converting all bimolecular uptake rates into their pseudo-first order equivalents:

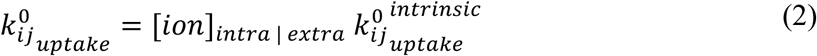

By folding in the effects of concentration as a pseudo-first order rate constant, we allow for a difference in chemical potential, μ, to drive flux. We measure the chemical potential created by a gradient in J/mol and refer to this interchangeably with concentration, e.g. an I-μ curve is a current-chemical potential or current-concentration curve. The Nernst equation allows us to define the chemical potential (concentration) gradient that exactly offsets an applied voltage:

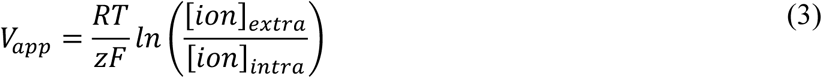

where R is the gas constant, T is temperature, z is the charge of the ion, F is Faraday’s constant, V_app_ is the applied voltage that would oppose the chemical gradient, [ion]_extra_ is the mM extracellular ion concentration, and [ion]_intra_ is the mM intracellular ion concentration. When we refer to equivalent voltage (V_eq_), we use this equation to calculate the voltage, but since V_eq_ is meant to drive flux in the same direction as the chemical potential, we use the negative of the applied voltage (V_app_). Thus, V_eq_ enables a direct comparison between the ion flux driven by voltage and ion flux driven by chemical potential.

For systems with sequential binding sites, the steady state flux through the channel can be calculated at any given transition. For example, in a channel with two binding sites flux can be calculated in three separate ways, one for each transition state region:

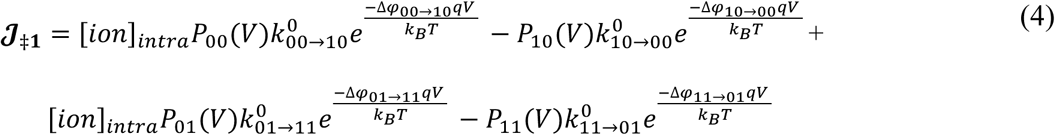

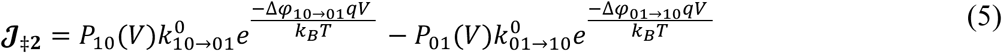

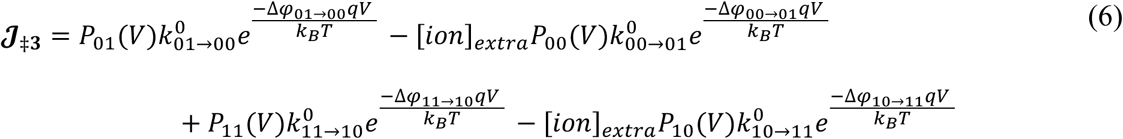

where 𝓙_‡**1**/**2**/**3**_ are the fluxes between intracellular bulk and binding site 1, site 1 and site 2, and site 3 and extracellular bulk, respectively; [ion]_extra_ is the mM extracellular ion concentration; and [ion]_intra_ is the mM intracellular ion concentration. Under steady state conditions, each of these three fluxes are equal, permitting any of the three equations to be selected for flux analysis of the model. However, the direct influence of both the voltage coupling and the intracellular and extracellular ion concentrations is unique in each equation.

### 2.2 Model Construction

Two different thermodynamic forces can drive the transport of charged substrates across the membrane: voltage, mediated by an electric field, and chemical potential, mediated by bulk substrate concentrations. To clearly illustrate the effects of these gradients, we establish models with two, three, or four stable binding sites. The networks for 2-site and 3-site models are shown in Figure 1. In addition to site count, these models have one of three symmetries**: perfectly symmetric**, in which equally spaced binding sites have identical transition rates and voltage sensitivities (Δ*φ* values); “**mirror symmetric**” (Figure 2), in which binding/release/transfer parameters can vary but do so equivalently for inward and outward flux; or **asymmetric**, in which transition rates and their voltage sensitivities are allowed to be unequal for inward and outward flux, inducing rectification. When rectification is induced, the magnitude of rectification is quantified as a rectification ratio (RR):

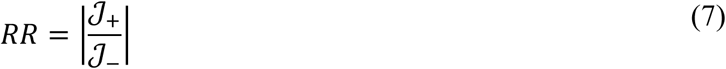

**Figure 1.**
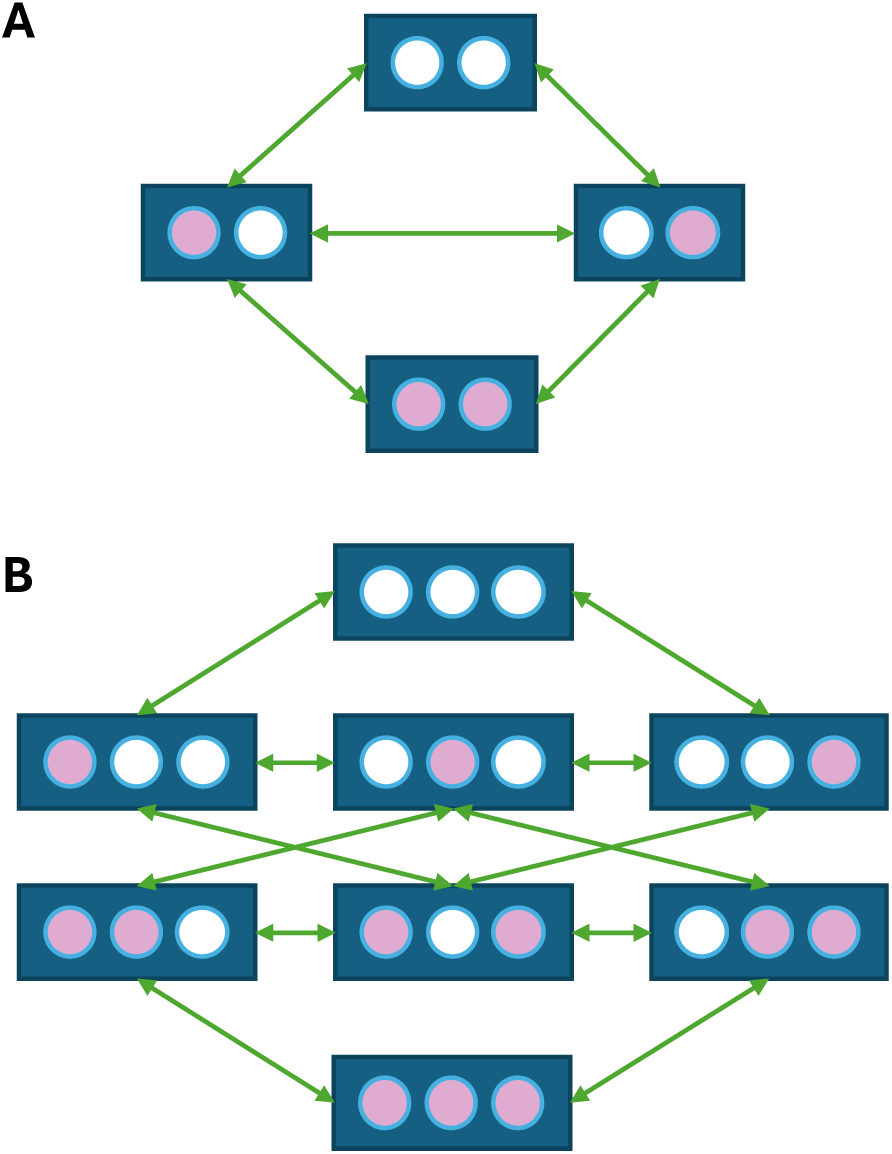
Intermediates states (nodes) and transitions (edges) highlighting kinetic network connectivity for channels with (A) 2 binding sites (2-site model) and (B) 3 binding sites (3-site model).

**Figure 2.**
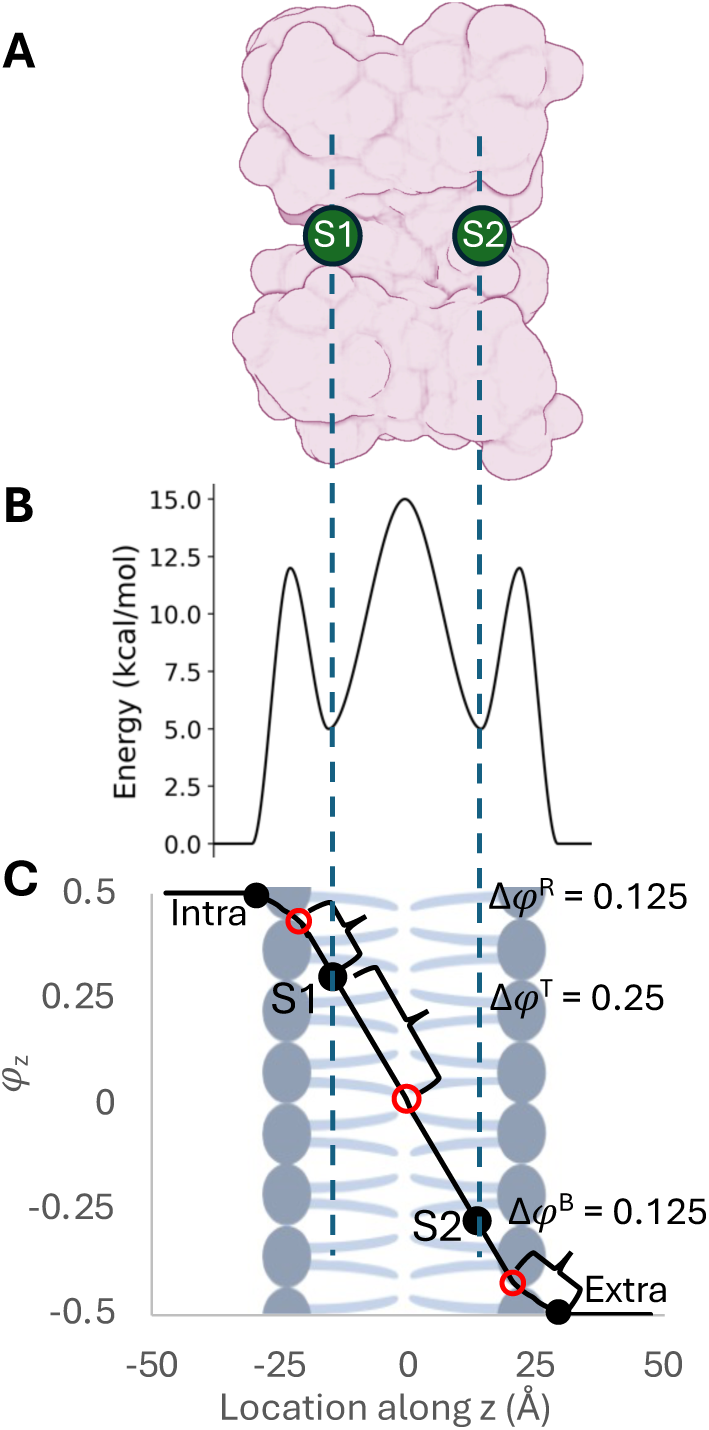
Two types of mirror symmetric systems. (A) Protein with two binding sites. (B) Rate mirror symmetry with different binding, release, and transfer barriers that are the same in the two directions. (C) Mirror symmetry of voltage sensitivity showing i1cp values for inward flux. Black filled circles represent binding sites and red circles represent transition states.

where 𝒥_+_ is the flux at a positive voltage and 𝒥_-_ is the flux at a negative voltage of the same magnitude for voltage rectification. For chemical rectification, the same definition holds except we use equivalent voltages. For both types of rectification ratio, the net chemical concentration ([ion]_extra_ + [ion]_intra_) is held constant between the positive and negative flux measurements. This definition means that outward rectifying systems have RR > 1 and inward rectifying systems have RR < 1. Thus, for comparison between the two in terms of magnitude, the reciprocal of an RR may be taken.

For both 2-site and 3-site models, the base transition rates, which are defined as transitions involving states with either single ion occupancy (transfer and release) or no ion occupancy (binding), and the voltage couplings were set on a model by model basis as detailed in the SI. All other transition rates were computed by modifying these base values using electrostatic potential differences as described in previous work^16,22^ (SI Section S1.2). All models used a simple linear dimensionless coupling profile (Figure 2B) and, unless otherwise noted, used a set effective dielectric screening constant (*ε*^′^) of 200.

Note this effective dielectric constant is otherwise an independent parameter in the solution optimization, meant to capture the full screening effects of the channel on ion-ion repulsions in a site- and transition-specific manner. It was set to a consistent large value (200) in the simple models to minimize rate asymmetry introduced by electrostatic repulsion and permit ion loading of channels congruent with experimental and simulation observations. To isolate the effects of voltage sensitivity from electrostatic or free energy differences, the binding site positions were fixed at their symmetric locations for all electrostatic calculations.

Finally, previously developed 4-site models^16^ of the Shaker K_v_ channel were used to evaluate whether the trends observed in the simpler systems extend to more complex and realistic protein models. These models exhibit considerable asymmetry in site locations and rates. They also have a wide range of independently optimized transition-specific effective dielectric constants (*ε*^′^) meant to capture the complex channel-ion interactions that collectively screen ion-ion repulsions. Although these solutions were optimized to reproduce experimental Shaker K_v_ I-V data, they were developed for the purpose of testing the limits of top-down kinetic modeling. Thus, they are simplified representations of the real Shaker system with only four sites, no intervening waters explicitly included, and a model dimensionless coupling profile, as fully described in Weckel-Dahman et al.^16^ Two solutions were selected from this work as representative protein models, designated solutions 9 and 12. While both solutions are consistent with available experimental data,^41^ they show markedly different responses to voltage and chemical gradients not included in solution identification. Full model specifications are provided in the Supporting Information (SI Section S1.3).

## 3. RESULTS AND DISCUSSION

### 3.1 Conduction Profiles Reveal Flux Bottlenecks

Across all model systems (SI Section S1.3), we consistently observe that the shape of voltage-dependent conductance curves can be classified into one of three distinct profiles (Figure 3). Each conductance profile depends on the flux-limiting step (FLS, the transition that constrains ion flux the most under a given electrochemical gradient) and has unique defining mechanistic characteristics. The FLS is caused by either slow transition rates producing a rate-limiting step (RLS) or transitions with low Δφ values producing a voltage-limiting step (VLS). We describe these conductance shapes below and outline their distinguishing mechanistic features.

**Figure 3.**
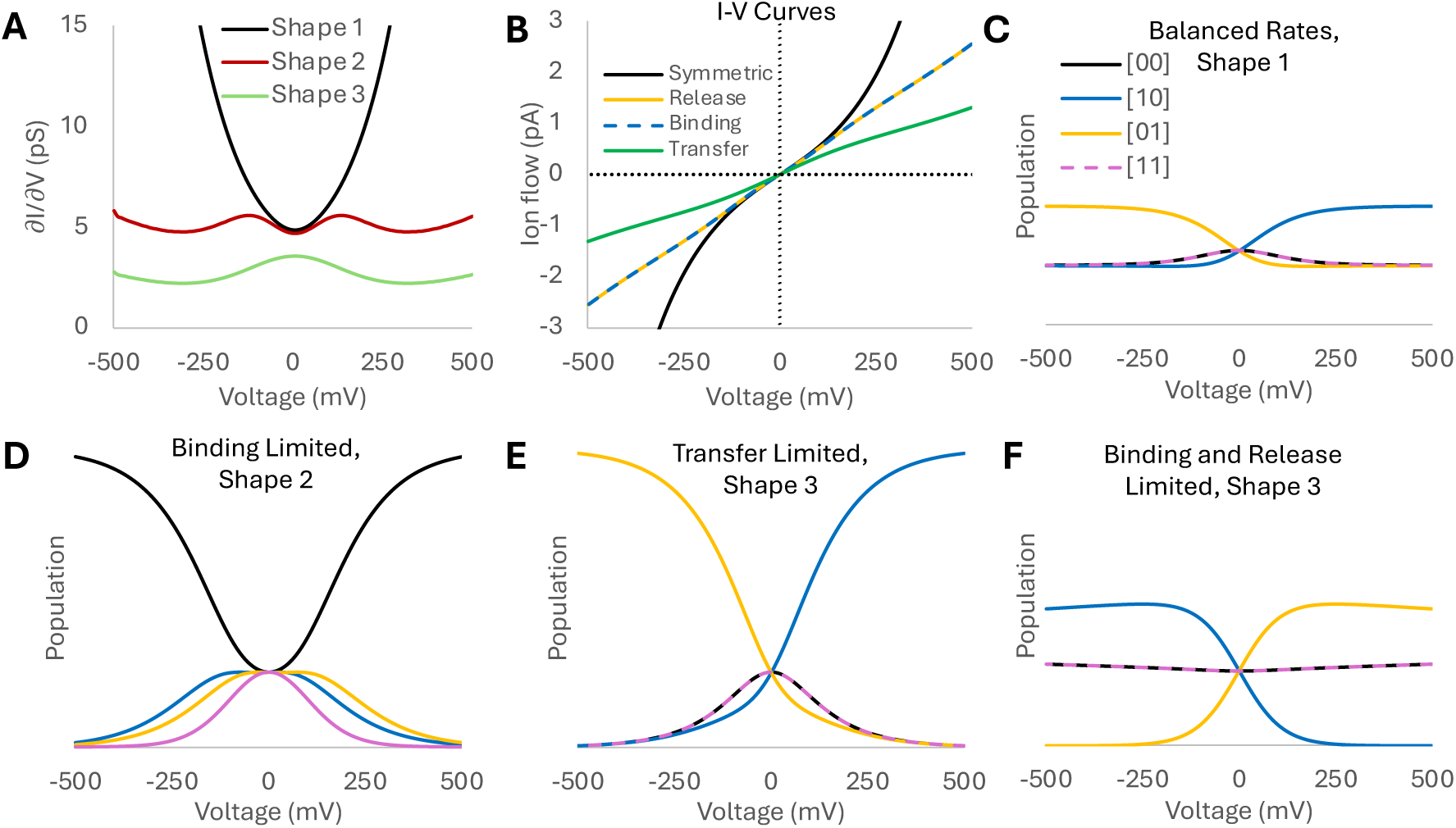
(A) Conductance curves, (B) I-V curves, and (C-F) corresponding populations for the three distinct conductance shapes. Conductance shape 1 (black) is the result of relatively equal fluxes resulting in increasing partial occupancy populations (C). Conductance shape 2 (red) reflects either a binding FLS (D) or a release FLS. Conductance shape 3 (green) reflects either a transfer FLS (E) or simultaneous binding and release FLSs (F). All population plots are equally scaled (y axis between 0 and 1).

**Conductance shape 1 (Balanced,** Figure 2B black) has a central minimum as its only extremum. Mechanisms associated with this shape have very similar flux through all transitions, resulting in a FLS that is only slightly slower than other transitions. The magnitude and location of the FLS is voltage dependent, switching from transfer-limited at experimentally relevant voltages to binding-limited at extreme voltages. Population analysis reveals that as voltage is increased, the fully ion-occupied state ([11]) and empty protein state ([00]) populations both decrease as they are partially converted into states [10] and [01] for positive and negative voltages, respectively. However, all states retain population. Similar but more extreme behavior is observed for transfer-limited conductance shape 3 (Figure 2E) with [11] and [00] dropping to negligible populations above 500 mV.

**Conductance shape 2 (Binding/Release Limited,** Figure 2B red) occurs in systems with large flux asymmetries between binding and release. A core characteristic of conductance shape 2 is that the FLS (either on binding or release) does not change with increasing voltage. Mechanisms associated with this conductance shape have dominant pathways with a voltage-dependent build-up of either the [00] state when binding limited (Figure 2D) or the [11] state when release limited. In these scenarios, ions initially move rapidly out of or into the channel, decreasing the populations of [11] (when binding-limited) or [00] (when release limited) while slightly increasing partial state occupancies (seen in the peak in conductance at ∼±175 mV for the system shown in Figure 2B). Beyond this point, these partial state occupancies decrease, resulting in non-zero minima in the conductance (∼±300 mV for the system shown in Figure 2B) (SI Figure S1). Eventually, the exponentiated decrease in the free energy barrier due to increasing voltage results in increasing flux.

**Conductance shape 3 (Transfer Limited** or **Binding + Release Limited**, Figure 2B green) occurs when transfer is flux limiting or binding and release are ∼equally flux limiting while transfer is fast. This shape is marked by a central peak at 0 mV. For the transfer limited case (Figure 2E), initially equal binding site populations rapidly diverge as ions bind to the first binding site until voltage is capable of overcoming the high barrier of transfer. For the binding and release limited case (Figure 2F), ions readily transfer between the binding sites, leading to a build-up of ions at the second binding site as release is limited. Furthermore, slow binding flux results in minimal changes in the populations of [00] and [11] states as voltage is increased. Regardless of the limiting step, the dominant mechanistic pathways involve singly-occupied states ([01] or [10]) and a FLS that does not change with increasing voltage.

### 3.2 Voltage and Rate Contributions to the FLS

While each conductance shape is defined by a distinct FLS, the FLS itself can evolve over a range of electrochemical conditions. This is because the FLS is a consequence of the balance between rate- and voltage-limiting steps. Variations in relative rates and Δ*φ* values modulate this balance and influence conductance behavior.

Voltage sensitivity, which appears as Δ*φ* in the exponential terms of the generalized exponential sum (Equation 1), can generate a FLS by limiting the exponential increase of some rates more than others as voltage is applied. Starting from a perfectly symmetric 2-site model, we compared mirror model variants in which the Δ*φ* of binding, transfer, or release were limited, normalizing net Δ*φ* values across the membrane to 1 (SI Figure S2). These comparisons revealed that any deviation from perfect voltage-sensitivity symmetry reduced voltage-driven flux and resulted in alterations of the conductance shape congruent with section 3.1.

Alternatively, the FLS can be a consequence of slow transition rates. Higher transition state barriers between intermediates and/or infrequent attempt frequences result in slower rates, restricting ion flux through the associated transitions. These bottlenecks lead to imbalanced flux, producing the same conductance shapes discussed above (SI Figure S3).

When we combined VLS and RLS, the two limiting factors reinforced or cancelled each other in a predictable manner, e.g. a slow rate increases the limiting effect of low voltage sensitivity on the same step, while a fast rate can decrease it (SI Figure S4). We additionally note that when rates and voltage sensitivity both contribute to the FLS, voltage sensitivity has a larger effect on the FLS at higher voltages, while rate sensitivity has a larger effect at lower voltages. The crossover point determining which (RLS or VLS) is the primary driver of the FLS can be located using rectification ratios (RR), as discussed in section 3.4.

### 3.3 Chemically Induced Ion Flux

In this section, we explore chemically driven ion flux using a 3-site mirror symmetric model. Without applied voltage, our flux equation (Equation 1) transforms from a generalized exponential sum to a weighted summation of the forward and reverse fluxes. Thus, only the voltage-independent free energy surface and associated transition rates influence the net flux. To explore their relative impact, we varied the rates of binding, release, and transfer to be between 0.01x (very rate-limiting) to 100x (very fast) relative to the values of the other rate constants (Figure 4).

**Figure 4.**
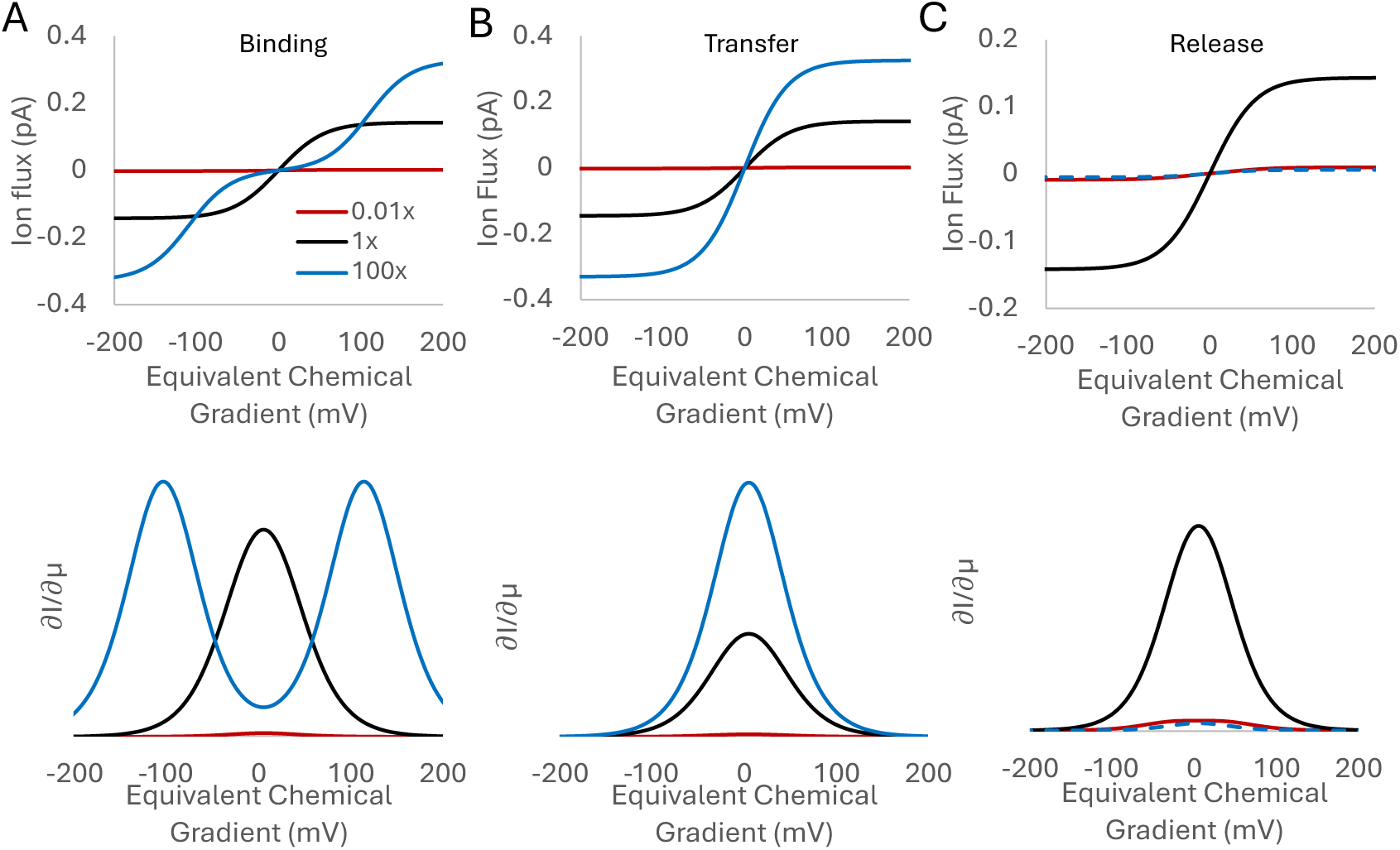
The influence of (A) binding, (B) transfer, and (C) release rate constants on concentration-driven (I-μ) flux (top) and chemical conductance (bottom). The black line shows the flux of a perfectly rate-symmetric system. 0.01x, 1x, and 100x refer to the scaling of the named rates relative to the other rates. Note the scale of release (C) is magnified to capture the subtle difference between 0.01x and 100x.

The binding rate constant in particular plays a significant role in the shape of I-μ curves (Figure 4A). When gradually increasing the binding rate constant from being very rate limiting (0.01x, Figure 4A red) to being equal (1x, Figure 4A black), we observe a logistic shape as previously reported.^22^ In this region, the channel is shifting from being mostly empty with limited flux due to slow uptake, to increasing flux with equally occupied sites responding to the chemical gradient (SI Figures S5). Past this point (Figure 4A blue), the FLS switches to release, reflecting the increasing stability of the binding sites relative to bulk such that reverse-uptake limits forward flux at small gradients until the gradient is large enough for release to again compete with reverse uptake enabling again forward flux (SI Figures S5).^22^ The curve shape also changes. Taking the derivative of these I-μ curves, which we will call chemical conductance, we observe a transition from one maximum to two maxima. These two maxima are increasingly separated as the binding rate is increased, reflecting the larger gradient needed to balance the now flux limiting release with reverse uptake. This is distinct from the trends seen when transfer and release are altered (Figure 4 B, C), where increasing or decreasing transfer or release rate constants do not alter the underlying shapes of the I-μ curves, but the relative flux of limiting (0.01x) and fast (100x) are clearly distinct between transfer and release. Again, explanations for these trends lies in the balance between forward and reverse flux in response to the chemical gradients. The full details of these trends and potential to extract relative rates directly from I-μ curves will be investigated in future work.

### 3.4 Understanding Current Rectification

All model systems analyzed to this point have been reflectively symmetric about the membrane normal (mirror symmetric). As soon as this symmetry is broken, an ion channel will exhibit rectification: preferential flux on one side of the IV curve. Focusing on open channel flux, rectification has two primary origins (excluding voltage-induced conformational changes or blockage): a free energy surface that is asymmetric about the membrane normal (rate-induced rectification) or asymmetric voltage sensitivity between transitions (voltage sensitivity-induced rectification). Classically, rectification has only been observed on I-V curves, but we will also explore rectification for chemical gradients, which we will call chemical rectification. This allows us to use I-V and I-μ curves to determine the primary cause of rectification in open channel current.

#### 3.4.1 Rate Induced Rectification

Starting with a perfectly symmetric 2-site model, rate asymmetry can be introduced either by changing the stability of a binding site or by changing the height of a transition state. As an example, asymmetry was introduced by making the S1 binding site 2 kcal/mol more stable than the S2 binding site. While this change led to minimal voltage rectification (RR of 1.0006 at ± 150 mV), we observed significant outward chemical rectification (RR of 1.428 at ± 150 mV, Figure 5). Generally, when rectification is induced via rate asymmetry, we observe that chemical rectification (I-μ) is more extreme than voltage rectification (I-V). When making a binding site more stable, the rectification mechanism depends on competition between uptake and transfer following ion release as previously observed.^16^ For proteins with low internal effective dielectric constants, resulting in higher ion-ion repulsion, this competition can be removed as ion-ion repulsions prevent reverse uptake from outcompeting forward transfer.^22^ However, we have found that increasing bulk concentration eventually ameliorates this repulsion and reintroduces reverse uptake competition. Thus, rectification that is more pronounced in I-μ curves and rectification that increases in magnitude with increasing total ion concentration are both signs of rate-induced rectification.

**Figure 5.**
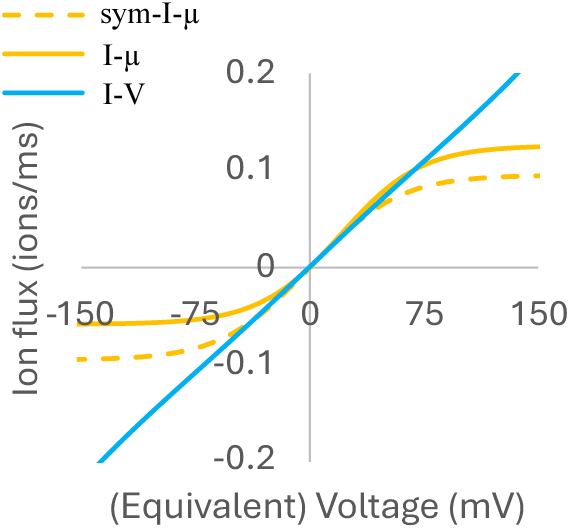
Rectification due to rate asymmetry when the stability of S1 is increased by 2 kcal/mol (solid) relative to symmetric rates (dashed). Rectification is apparent in I-μ curves (yellow), but hidden in I-V curves where symmetric and asymmetric overlap, indicating negligible voltage rectification. x-axis is voltage for I-V curves and equivalent voltage for I-m curves.

#### 3.4.2 Voltage-Sensitivity Induced Rectification

Voltage-sensitivity induced rectification can be caused by asymmetric binding site locations or when transition states between binding sites are asymmetrically placed. These two influences lead to different rectification patterns. Regardless, when rectification is induced via imbalances in voltage sensitivity (Δ*φ*), non-unitary RR(s) are only observed in I-V curves—not in the corresponding I-μ curves. Another general property of voltage rectification is that the RR increases as the voltage increases. Thus, eventually any degree of voltage asymmetry will lead to a larger voltage RR than the chemical RR. The crossover point where the voltage RR (measured in the I-V curve) becomes larger than the chemical RR (in the I-μ curve) is also where the VLS become the dominant influence on the system FLS, as mentioned in section 3.2.1.

##### 3.4.2.1 Voltage-Induced Rectification due to Binding Site Locations

The relative voltage sensitivity of every step depends on the distances between binding sites and transition states. Moving binding sites further apart while keeping the transition state placed equidistant between sites thus increases voltage sensitivity in equal but opposite directions for forward and reverse flux. For example, starting with a perfectly symmetric 2-site model, we gradually induce rectification by moving the extracellular binding site towards the intracellular bulk (Figure 6). As the extracellular binding site is moved, we find that the FLS becomes transfer-limited between sites S1 and S2 while uptake from and release to the extracellular solution become most voltage sensitive. The direction of rectification depends on bulk concentrations (shifting the balance in uptake and release), but regardless of the site or direction of movement, increased movement of a site leads to increased asymmetry in voltage sensitivities and hence increased RR.

**Figure 6.**
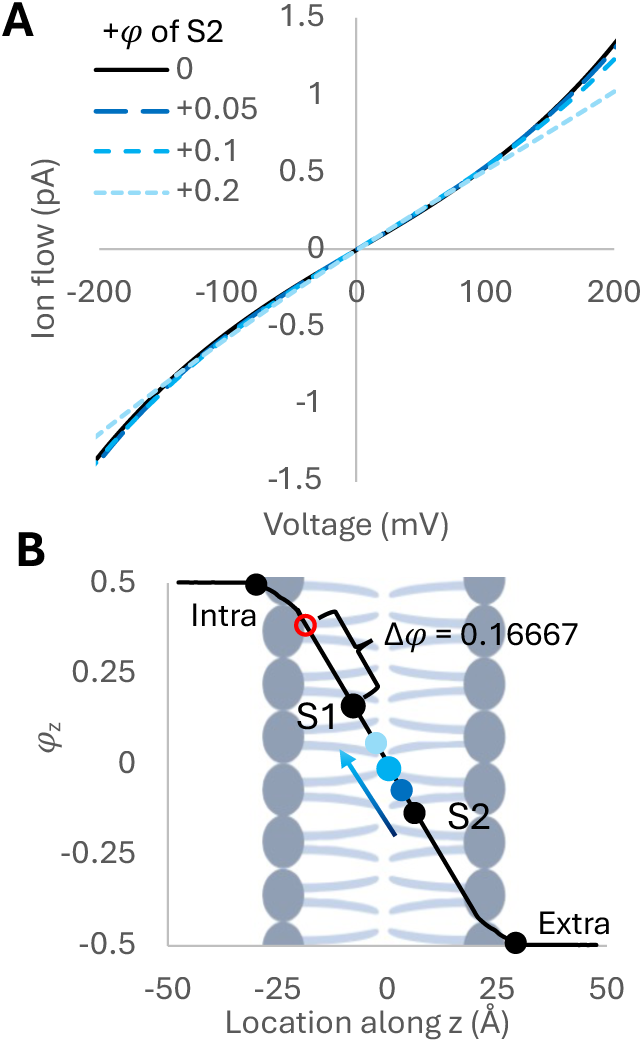
(A) Voltage-induced rectification due to moving extracellular binding site (S2) towards intracellular side, increasing voltage coupling for extracellular binding and release while decreasing it for transfer from S1 to S2. (B) Representative site and Δφ locations depicted on model membrane with the S2 location colors matching the colors of the resulting I-V curves

##### 3.4.2.2 Voltage-Induced Rectification due to Transition State Locations

When a transition state is not equidistant between two binding sites, the voltage-induced rectification is generally more pronounced. Starting from the same perfectly symmetric 2-site model, voltage asymmetry was introduced by shifting the location of the transition state between S2 and the extracellular bulk towards S2 (Figure 7), leading to stronger voltage coupling for binding to S2 and weaker voltage coupling for release from S2.

**Figure 7.**
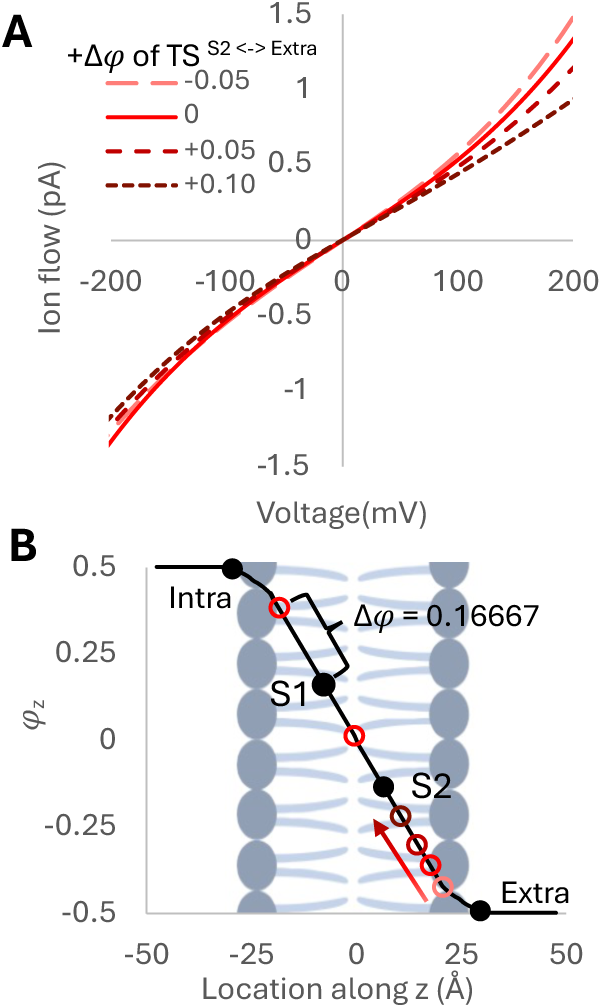
(A) Voltage-induced rectification due to moving transition state between S2 and extracellular bulk towards S2, progressively increasing voltage coupling of binding. (B) Representative transition state depicted on model membrane with colors matching the colors of the resulting I-V curves in (A).

Since moving a transition state causes unequal sums of inward and outward Δ*φ* values, we find that transition state shifts generate stronger voltage-induced rectification. At ±150 mV, moving the *φ* of S2 +0.1 closer to S1 results in a RR of 0.928, compared to an RR of 0.842 for a +0.1 *φ* shift in the transition state between S2 and extracellular bulk. Interestingly, the ocation of the relative voltage sensitivities (binding/transfer/release) is not as important as their relative magnitudes in inward and outward flux. We found equivalent I-V curves when transition state shifts occur in the same flux direction (SI Figure S6) for different transition types that give the same conductance shape. For example, equivalent rectifying I-V curves can be found when setting the |Δ*φ*| of binding at S1 to be 0.1 or the |Δ*φ*| of release at S2 to be 0.1 (SI Figure S6, Model 1, A and B). Thus, asymmetry in the location of the transition state relative to binding sites in the forward and reverse directions introduces strong voltage-induced rectification that manifests in larger RR(s) in I-V curves relative to RR(s) observed from shifting binding site locations.

### 3.5 Example Protein Systems

Real channels are of course much more complicated than the model systems studied above. It is expected that they will have competing and/or complimentary asymmetry in both rates and voltage sensitivity, in addition to site and even transition-specific ion-ion repulsions balanced by channel screening. To test whether the observations from model systems still hold when these more complex facets influence open-channel conductance, we used two kinetic solutions previously identified for the Shaker K_v_ channel.^16^ This was deemed essential since there are no known complete kinetic solutions (known transitions and transition rates) for a real channel. Since the rate matrices are known exactly for these solutions, this enables us to interpret trends based on known rate and voltage asymmetries.

We selected two solutions (solutions 9 and 12) that have unique sets of rates but nearly identical mechanisms, with flux differences arising only in nondominant pathways that collectively contribute <1% of the net flux. Although these two solutions produce practically identical mechanisms and flux for the voltages included in their original solution optimization, they produce dramatically different flux for conditions not included in their optimization—such as larger voltages (SI Figure S7) and chemically-driven flux (Figure 7). The similarity between these two solutions under certain conditions, combined with differences under other conditions, provides an informative test of the principles discussed in this paper.

#### 3.5.1 Looking for the Origin of Channel Rectification

Initially, we wanted to assess which rectification mechanism is dominant in solutions 9 and 12. Since both solutions have identical voltage sensitivities by design, rate differences must be the cause of their different electrochemical responses. But it is not known which is more pronounced—rate or voltage asymmetry. This can be extracted from electrophysiology assays by comparing RR(s) in I-V and I-μ curves, as discussed in section 3.4.

First, we generated I-V and I-μ curves between ±150 mV with 200 mM net K^+^ ions available, which is outside of the range for which these models were optimized. Next, we used the RR assessed at ±150 mV to compare the relative impact of voltage and rate asymmetry. Both solutions show larger asymmetry in their I-μ curves (Figure 8), indicative of larger rate asymmetries. Solution 9 has an inward RR of 0.36 in I-μ versus only 0.56 in I-V. For solution 12, the difference is even more pronounced, with opposite directions for chemical and voltage rectification.

**Figure 8.**
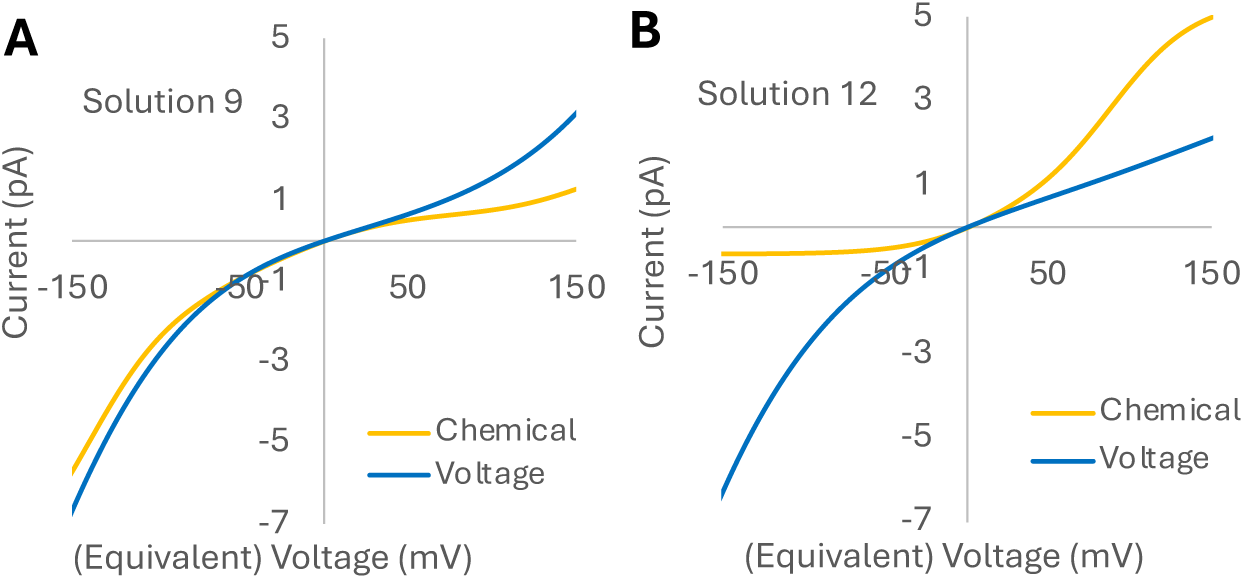
I-V (blue) and I-μ (yellow) curves for (A) solution 9 and (B) solution 12.

Chemical rectification for solution 12 was strongly outward rectifying (RR of 4.96 outward, with a reciprocal of 0.2 for purposes of comparison with solution 9’s inward rectification) versus inward rectification of 0.27 in I-V. The larger and flipped I-μ rectification of solution 12 suggests that its rate asymmetry should be larger than solution 9’s, which it is, and that the direction of the rate asymmetry should also flip to oppose the direction of the voltage sensitivity, which it does. Even when comparing between solutions, the differences in I-V RR(s) are less significant than differences in I-μ RR(s), consistent with our discussion of rates having a smaller impact on the I-V rectification in section 3.4.1.

In summary, comparing relative RR(s) in I–V versus I–μ profiles correctly identifies whether rectification arises from voltage sensitivity or from rate asymmetry, both within a single system and when comparing systems. This assignment is voltage-specific: as the underlying mechanistic origin changes with voltage, the RR trend changes accordingly. It is also direction-specific, with rectification for inward versus outward flux manifesting in RR(s) at negative versus positive voltages (for cations), respectively, such that the flux in either direction can be independently interpreted.

#### 3.5.2 Extracting the FLS from Conductance Profiles

Next, we sought to learn which steps are flux limiting based on conductance curves. Starting with solution 9, both sides of the I-V curve follow conductance shape 1 up to ±90 mV (Figure 9A). Thus, we expect to see relatively balanced FLS(s). Mechanistic analysis of the dominant flux pathways in each direction for solution 9 is consistent with conductance shape 1, with the dominant inward and outward pathways containing nearly identical flux through all transitions. These findings reinforce that the distribution and symmetry of the FLS(s) in the dominant flux-producing pathways govern the resulting conductance shape.

**Figure 9.**
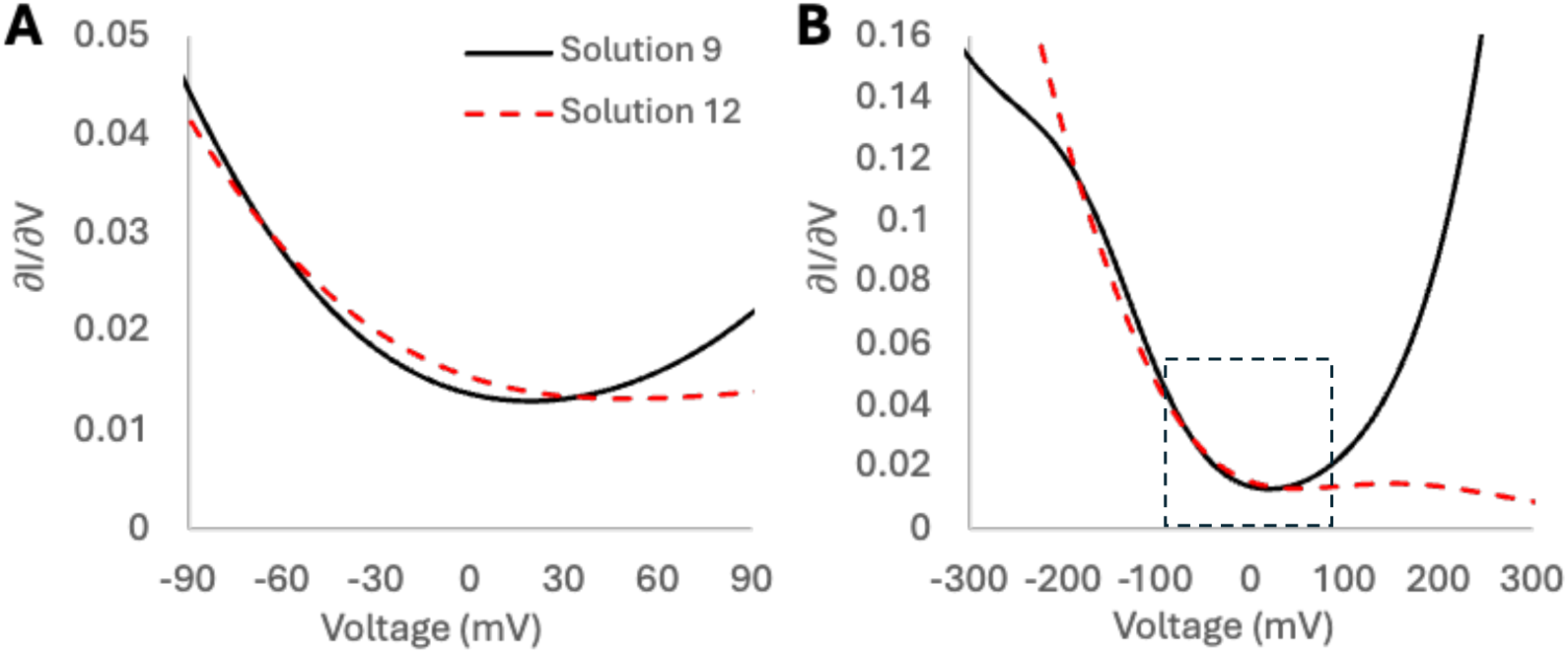
Conductance of the ion flux (Figure 8 blue) for solution 9 (black) and solution 12 (red dashed) at (A) low magnitudes of the voltage and (B) an extended voltage range.

Similar to solution 9, solution 12 is expected to have a balanced FLS at -90 mV, while the marked decline in flux at +90 mV indicates mechanism changes. The lack of a central peak in the conductance suggests that solution 12 transitions to conductance shape 2 for positive voltages. This is clearer as voltage is taken to larger values with a slight increase up to +150 mV followed by a decrease (Figure 9B) tracking shape 2 (Figure 3B, red). Detailed mechanistic analysis confirms these predictions. At -90 mV, the top two flux producing cycles are consistent with conductance shape 1 with multiple FLS(s) of similar magnitude and symmetric flux distributions on the negative voltage side (inward flux for cations). At +90 mV we observe a single binding-limited dominant pathway (∼56.7% of the total flux) with significant binding/release flux asymmetry (specifically binding limited in this case), consistent with conductance shape 2. Notably, for solution 9, we find that the FLS of the dominant pathway shifts around -200 mV that also has binding/release asymmetry consistent with shape 2. For both systems, the conductance profiles correctly indicate where flux is limiting—and they do so independently for the two flux orientations.

## CONCLUSIONS

The goal of this study was to understand the fundamental physical principles that govern an open ion channel’s response to changing electrochemical conditions. Focusing on open-channel conductance, we systematically altered protein properties that are expected to impact electrochemically induced ion flux. Specifically, we probed the influence of relative rates and voltage sensitivity for ion uptake, transfer, and release (with the voltage sensitivity being dictated by binding site and transition state locations) using two and three site electrochemically-responsive kinetic models.^16,23^ By analyzing I-V and I-μ curves across a range of electrochemical conditions, we are able to relate electrophysiology assays to the physical and mechanistic properties that cause non-ohmic current and rectification. For voltage-driven flux, three conductance profile shapes were found—each with distinct FLS(s) and preferential occupancy in their dominant pathways as a function of the applied voltage.

A central feature of this work is the value of evaluating not just voltage-driven flux curves (I-V), but also chemical driven (I-μ). In addition to being distinct from I-V curves, I-μ curves reveal when rectification is dominated by asymmetry in relative rates, manifesting in larger ‘chemical rectification’ ratios in I-μ curves than ‘voltage rectification’ ratios in I-V curves. In contrast, systems that are asymmetric in voltage sensitivity but not in rates will show more pronounced rectification in I-V curves. In this manner differences in flux between purely chemical and purely voltage gradients reveal the physical origin of open channel rectification. Additionally, chemical conductance profiles can reveal when a system is uptake limited by transitioning from a single to bimodal maxima. The full ramifications of this will be explored in future work.

Extending our analyses to more complex systems, we conduct flux and conductance analyses of two 4-site kinetic solutions for the Shaker K_v_ channel (solutions 9 and 12) as model ion channels with known mechanisms and underlying transition rates. We find that the predictions from simple model systems hold. Not only do voltage- vs. rate-induced rectification relationships manifest in larger rectification ratios in I-V vs. I-μ curves, respectively, but the relative orientation of asymmetry is extracted as well. Turning to conductance profiles, the underlying balance vs. asymmetry in which transitions are flux limiting is consistent with the three predicted conductance shapes in a direction-specific manner (meaning positive and negative flux profiles can be interpreted independently). We also find that conductance curves can transition from one conductance shape to another under changing voltage conditions when the flux-limiting step is also voltage-sensitive.

Together, these findings support the broader applicability of using I-V and I-μ curves to dissect how specific electrochemical and kinetic parameters shape open-channel behavior. Specifically, this framework offers an experimentally testable approach for inferring: 1) the location of the FLS, 2) the relative impact of rates and voltage-sensitivities on rectification, 3) the concentration at which binding becomes flux limiting, and 4) when changing electrochemical conditions causes the dominant mechanism to shift from a pathway with one set of these features to another. It also offers a quantitative bridge to compare mechanisms observed in simulations to experimental electrophysiology data. However, caution should be taken in applying this analysis to systems in which current-voltage relationships are strongly influenced by other factors, such as the titration-dependent build-up of ions at the channel entrance, voltage-sensitive lipid-interactions, or voltage-induced conformational changes, including but not limited to gating. Nevertheless, the presented framework describes how electrochemical gradients alone influence open-channel currents (i.e., before or after gating and in addition to the above influences). Since these effects are always present, they form a foundation that must be factored into more complex models that also account for titration, gating, and/or lipid-regulation. Such integrations and extending these methods to real channels will also be the focus of future work.

## ASSOCIATED CONTENT

### Supporting Information

The following files are available free of charge.

- Supplementary information file includes additional derivations (Sections S1.1 and S1.2), model specifications (Section S1.3), and results (Section S2). These results include separated population curves for conductance shape 2 (Figure S1), full scan of Δ*φ* values used to assess the impact of the VLS on conductance shape (Figure S2), full scan of k_ij_ values with 3-site protein models that used perfectly symmetric (Figure S3) and asymmetric (Figure S4) Δ*φ* values, PMF curves for the chemical conductance curves of Figure 3 (Figure S5), equivalent I-V and conductance curves in voltage-rectifying systems (Figure S6) with their respective voltage sensitivities used (Tables S5 and S6), and I-V curves for Solutions 9 and 12 (Figure S7).

## AUTHOR INFORMATION

### Author Contributions

H.W.D, R.C., and J.M.J.S designed the research. H.W.D., R.C., and A.D.D. built and analyzed the kinetic models. All authors helped interpret the results. J.M.J.S directed and funded the research. All authors wrote the manuscript and have given approval on the final version.

### Notes

The authors declare no competing financial interest.

## ACKNOWLEDGMENT

The authors would like to thank Professor Aurora Clark for helpful discussions. This work was supported by NIH NIGMS (R35GM143117) and the computational resources provided by Expanse at the San Diego Supercomputing Center through the Advanced Cyberinfrastructure Coordination Ecosystem: Services and Support (ACCESS) program (allocation MCB200018) supported by NSF (grant nos. 2138259, 2138286, 2138307, 2137603, and 2138296), as well as the Center for High-Performance Computing (CHPC) at the University of Utah.

## ABBREVIATIONS

I-V: current voltage
I–μ: current concentration
MD: molecular dynamics
Δ*φ*: voltage sensitivity
FLS: flux-limiting step
RLS: rate-limiting step
VLS: voltage-limiting step
[11]: fully ion-occupied state
[10]: intracellular binding site occupied state
[01]: extracellular binding site occupied state
[00]: empty protein state
RR: rectification ratio

## TOC Graphic

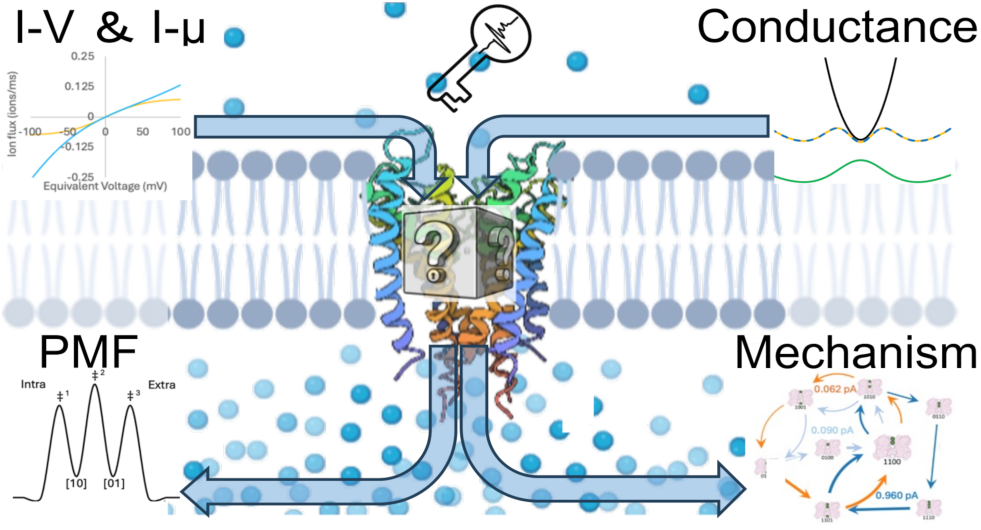

## Notes

### Competing Interest Statement

The authors have declared no competing interest.

