## Supplemental Information for "From Flux to Function: Extracting Mechanistic Insights from Ion Channels via I-V and I-μ Analyses"

### S1. SUPPLEMENTAL METHODS

#### S1.1 Derivation for Voltage-responsive Rate Constants and Biomolecular Flux

The 4-site Shaker solutions were calculated $\varphi(z)$ using the continuum electrostatic approximation method,[1,2] which takes the difference between ensemble-averaged electrostatic potentials ($\Delta\phi_{elec}^{\pm V}(z)$) for simulations run with a negative $\Delta\phi_{elec}^{-V}(z)$ and positive $\Delta\phi_{elec}^{+V}(z)$ voltage of equal magnitude and normalizes the expression with the total voltage drop, $V_T$:

$$\varphi(z) = \frac{\Delta\phi_{elec}^{-V}(z) - \Delta\phi_{elec}^{+V}(z)}{V_T} \tag{S1}$$

In kinetic modeling, rate constants of rare-event transitions are defined by the difference in free energy between meta-stable intermediates and transition barriers ($\Delta G_{ij}^{\ddagger}$). Using Roux's W-route,[3] we previously described the free energy difference under a transmembrane potential as:[4]

$$G(z,V) = \varphi(z)qV + G(z,0) \tag{S2}$$

$$\Delta G_{ij}^{\ddagger}(z,V) = \left[\varphi_{ij}(z^t)qV + G(z^t,0)\right] - \left[\varphi_{ij}(z^r)qV + G(z^r,0)\right] \tag{S3}$$

where q is the charge of the ion moving in the transition, V is the transmembrane voltage, and $G(z^t,0)$ and $G(z^r,0)$ represent the zero-voltage free energy at the transition barrier and reactant basin, respectively. Using the Eyring-Polanyi equation (Equation S4), we then derive a relationship between rate coefficients under different transmembrane potentials:[4]

$$k_{ij}^{0} = \frac{\omega_0}{2\pi} e^{\frac{-\Delta G_{ij}^{\ddagger}(z,0)}{k_B T}} \tag{S4}$$

$$k_{ij}(V) = k_{ij}^{0} e^{\frac{-[\varphi_{ij}(z^t)-\varphi_{ij}(z^r)]qV}{k_B T}} \tag{S5}$$

where $k_B$ is Boltzmann's constant, T is the temperature, $\omega_0$ is the attempt frequency, and $k_{ij}^{0}$ is the zero-voltage rate coefficient for a transition from state i to j.[4,5]

With voltage responsive rates, the flux associated with a rate is:

$$\mathcal{J}_{ij}(V) = P_i k_{ij}(V) \tag{S6}$$

where $\mathcal{J}_{ij}$ is the flux from state i to j and $P_i$ is the population of state i.

**S1.2 Relating Electrostatics to Rate Constants in MsRKM**

Since free energy and electric potential energy are additive, $\Delta G^{\ddagger}$ for each transition is a summation of the activation energy of the empty channel and the electric potential energy due to electrostatic attractive/repulsive forces between ions.

$$\Delta G_{ij}^{\ddagger} = \Delta G_{empty}^{\ddagger} + \Delta E_{elec} \tag{S7}$$

The electric potential energy $E_{elec}$ of two point charges $q_m$ and $q_n$ separated by a distance r is:

$$E_{elec} = \frac{1}{4\pi\varepsilon_0\varepsilon'}\frac{q_m q_n}{r_{mn}} \tag{S8}$$

where $\varepsilon_0$ is the permittivity of free space and $\varepsilon'$ represents the effective dielectric constant. The effective dielectric constant is a singular term that represents the dielectric constant of the solvent, as well as any local shielding effects that can modify the potential.[6] Similar to the free energy of a transition, this can be calculated for a particular transition as the net difference in electric potential energy experienced by an ion as it moves between a reactant basin and a transition state:

$$\Delta E_{elec} = E_{elec}^{T} - E_{elec}^{R} \tag{S9}$$

$$\Delta E_{elec} = \frac{1}{4\pi\varepsilon_0\varepsilon'} q_m q_n \left(\frac{1}{r_{mn}^{T}} - \frac{1}{r_{mn}^{R}}\right) \tag{S10}$$

With $r_{mn}^{T}$ and $r_{mn}^{R}$ being the distance between an ion at the transition state and reactant position for an ion at a binding site within the protein.

Combining equations S10 and S12 with equation S4 we get:

$$k_{ij} = \frac{\omega_0}{2\pi} e^{-\frac{\Delta G^{\ddagger}_{empty} + \Delta E_{elec}}{k_B T}} \tag{S11}$$

$$k_{ij} = k_{ij}^{Base} e^{-\frac{\Delta E_{elec}}{k_B T}} \tag{S12}$$

where $k_{ij}^{Base}$ is the transition rate from site i to j with an empty transporter.

### S1.3 Model Design

For all models in this section, we used a generic membrane thickness of 40 Å. Unless specified differently in the text below, binding site locations for electrostatic calculations were placed equidistant from each other and the membrane boundary. Assigned phi values assume a linear voltage drop, as in previous studies.[4,7,8] Transition states (TS) are located at the halfway distance between binding sites or between a binding site and the adjacent membrane boundary unless otherwise specified.

#### *S1.3.1 2-site Symmetric Model*

The 2-site perfectly symmetric model was designed to maximize the total symmetry in the system. Thus, $\Delta\varphi$ values are all equal and $k_{ij}^{Base}$ are equal. Bulk concentration was set to 1 mM, ensuring a 1:1 ratio for all rates at zero applied voltage. For the 2-site model, binding sites were placed at $\varphi$ values of $\pm \sim 0.1\overline{6}$, with a final $\Delta\varphi$ for each transition of $\pm\ 0.1\overline{6}$ and $\sim 13.\overline{3}$ Å apart from one another. Due to the distance between binding sites, rates modified by electrostatics were up to ~1% faster (for transitions that moved with electrostatic repulsion) or slower (for transitions that moved against electrostatic repulsion) than the initial base rates.

*S1.3.2 3-site Symmetric Model*

Like the 2-site model, the 3-site symmetric model was designed to maximize symmetry about $\Delta\varphi$ values and the final $k_{ij}$. For the 3-site model, this used binding sites at $\varphi$ values of $\pm$ ~0.250 and 0, with a final $\Delta\varphi$ for each transition of $\pm$ 0.125 with binding sites placed ~10 Å apart. Rates modified by electrostatics with the maximum ion occupancy were up to ~3% faster or slower than the initial base rates.

*S1.3.3 2-site and 3-site Mirror-symmetric Models*

In order to pull trends from changing $\Delta\varphi$ values, electrostatic effects were kept consistent with the perfectly symmetric 2-site and 3-site models. Models are presented in order as discussed in the main text.

Table S1: MsRKM model specifications used for all 2-site mirror symmetric models.

| | Model | System | $\lvert\Delta\varphi\rvert^{Bind}$ | $\lvert\Delta\varphi\rvert^{Release}$ | $\lvert\Delta\varphi\rvert^{Transfer}$ | $k_{ij}^{all}$ |
|---|---|---|---|---|---|---|
| Figure 3 | 2-site | Shape 1 | 0.166666 | 0.16 | 0.16 | 1 |
| | | Shape 2D | 0.05 | 0.225 | 0.225 | 1 |
| | | Shape 3E | 0.225 | 0.225 | 0.05 | 1 |
| | | Shape 3F | 0.05 | 0.05 | 0.4 | 1 |
| SI Figure S2A | | Dark Green | 0.245 | 0.01 | 0.245 | 1 |
| | | Green | 0.225 | 0.05 | 0.225 | 1 |
| | | Light Green | 0.2 | 0.1 | 0.2 | 1 |

| | | | | | | |
|---|---|---|---|---|---|---|
| | | Light Purple | 0.15 | 0.2 | 0.15 | 1 |
| | | Purple | 0.1 | 0.3 | 0.1 | 1 |
| | | Dark Purple | 0.05 | 0.4 | 0.05 | 1 |
| SI Figure S2B | | Dark Purple | 0.245 | 0.245 | 0.01 | 1 |
| | | Purple | 0.225 | 0.225 | 0.05 | 1 |
| | | Light Purple | 0.2 | 0.2 | 0.1 | 1 |
| | | Light Green | 0.15 | 0.15 | 0.2 | 1 |
| | | Green | 0.1 | 0.1 | 0.3 | 1 |
| | | Dark Green | 0.05 | 0.05 | 0.4 | 1 |

Table S2: MsRKM model specifications used for all 3-site mirror symmetric models.

| | Model | System | $k_{ij}^{Bind}$ | $k_{ij}^{release}$ | $k_{ij}^{Transfer}$ | $\|\Delta\varphi\|^{Release}$ | $\|\Delta\varphi\|^{B/T}$ |
|---|---|---|---|---|---|---|---|
| SI Figure S3A | 3-site | Dark Red | 0.01 | 1 | 1 | 0.125 | 0.125 |
| | | Red | 0.1 | 1 | 1 | 0.125 | 0.125 |
| | | Black | 1 | 1 | 1 | 0.125 | 0.125 |
| | | Blue | 10 | 1 | 1 | 0.125 | 0.125 |
| | | Dark Blue | 100 | 1 | 1 | 0.125 | 0.125 |
| SI Figure S3B | | Dark Red | 1 | 0.01 | 1 | 0.125 | 0.125 |
| | | Red | 1 | 0.1 | 1 | 0.125 | 0.125 |
| | | Black | 1 | 1 | 1 | 0.125 | 0.125 |
| | | Blue | 1 | 10 | 1 | 0.125 | 0.125 |
| | | Dark Blue | 1 | 100 | 1 | 0.125 | 0.125 |
| SI Figure S3C | | Dark Red | 1 | 1 | 0.01 | 0.125 | 0.125 |
| | | Red | 1 | 1 | 0.1 | 0.125 | 0.125 |
| | | Black | 1 | 1 | 1 | 0.125 | 0.125 |
| | | Blue | 1 | 1 | 10 | 0.125 | 0.125 |
| | | Dark Blue | 1 | 1 | 100 | 0.125 | 0.125 |
| Figure 4A | | Red | 0.01 | 1 | 1 | 0.125 | 0.125 |
| | | Black | 1 | 1 | 1 | 0.125 | 0.125 |
| | | Blue | 100 | 1 | 1 | 0.125 | 0.125 |
| Figure 4B | | Red | 1 | 1 | 0.01 | 0.125 | 0.125 |
| | | Black | 1 | 1 | 1 | 0.125 | 0.125 |
| | | Blue | 1 | 1 | 100 | 0.125 | 0.125 |

| | | | | | | | |
|---|---|---|---|---|---|---|---|
| Figure 4C | | Red | 1 | 0.01 | 1 | 0.125 | 0.125 |
| | | Black | 1 | 1 | 1 | 0.125 | 0.125 |
| | | Blue | 1 | 100 | 1 | 0.125 | 0.125 |

*S1.3.4 Asymmetric Models*

In order to pull trends from changing $\Delta\varphi$ values, electrostatic effects were kept consistent with the perfectly symmetric 2-site model. Models are presented in order as discussed in the main text.

Table S3: Model specifications used for the 2-site asymmetric models.

| | | $k_{ij}$ | | | | | | $\|\Delta\varphi\|$ | | | | | |
|---|---|---|---|---|---|---|---|---|---|---|---|---|---|
| | System | Intra -> S1 | S1 -> Intra | S1 -> S2 | S2 -> S1 | S2 -> Extra | Extra -> S2 | Intra -> S1 | S1 -> Intra | S1 -> S2 | S2 -> S1 | S2 -> Extra | Extra -> S2 |
| Figure 4 | All | 1 | 10 | 10 | 1 | 1 | 1 | 0.16 | 0.16 | 0.16 | 0.16 | 0.16 | 0.16 |
| Figure 5 | 0 | 1 | 1 | 1 | 1 | 1 | 1 | 0.16 | 0.16 | 0.16 | 0.16 | 0.16 | 0.16 |
| | +0.05 | 1 | 1 | 1 | 1 | 1 | 1 | 0.16 | 0.16 | 0.14167 | 0.14167 | 0.191665 | 0.191665 |
| | +0.1 | 1 | 1 | 1 | 1 | 1 | 1 | 0.16 | 0.16 | 0.11667 | 0.11667 | 0.216665 | 0.216665 |
| | +0.2 | 1 | 1 | 1 | 1 | 1 | 1 | 0.16 | 0.16 | 0.06667 | 0.06667 | 0.2666665 | 0.2666665 |
| Figure 6 | -0.05 | 1 | 1 | 1 | 1 | 1 | 1 | 0.16 | 0.16 | 0.16 | 0.16 | 0.216665 | 0.116665 |
| | 0 | 1 | 1 | 1 | 1 | 1 | 1 | 0.16 | 0.16 | 0.16 | 0.16 | 0.16 | 0.16 |
| | +0.05 | 1 | 1 | 1 | 1 | 1 | 1 | 0.16 | 0.16 | 0.16 | 0.16 | 0.116665 | 0.216665 |
| | +0.1 | 1 | 1 | 1 | 1 | 1 | 1 | 0.16 | 0.16 | 0.16 | 0.16 | 0.066665 | 0.266665 |

The $k_{ij}$ and $|\Delta\varphi|$ values are explicitly defined for each transition.

*S1.3.5 Solutions 9 and 12*

These more complex models were used to test the trends found in the simpler systems above. The models were previously developed to test the impact of including ion repulsions to reduce the kinetic solution space and the importance of retaining the full network as electrochemical gradients vary.[8] They were fit to Shaker $K_v$ channel data but included only four $K^+$ binding sites, one located in the gate region[9] (G1) and three located within the selectivity filter at T441 (S4), G443 (S2), and G445 (S0).[10,11] These site locations were selected from locations with high ion occupancy in molecular dynamics simulations of Shaker performed by Naranjo et. al.[12] These models focused on open channel currents, excluding gating c-type inactivation.[13] The models did not include perfectly coupled (i.e., simultaneous) transitions or transitions where an ion skips an intermediate site, which reduces the number of possible transitions to 56. However, base rates (those for single ion occupancy) were allowed to go as high as 1e13 $s^{-1}$, with even faster rates due to ion-ion repulsions, resulting in rates on the order of 1e14 $s^{-1}$, which is effectively simultaneous. For electrostatic and voltage-coupling calculations, the binding site and transition state locations in Table S1 were used with the simple linear voltage-drop approximation:

Table S4: Site locations used for solution 9 and solution 12. Membrane edges (not included in table) are located at ±20 Å.

| Model | Intra | TS 1 | G1 | TS 2 | S4 | TS 3 | S2 | TS 4 | S0 | TS 5 | Extra |
|---|---|---|---|---|---|---|---|---|---|---|---|
| Solutions 9 and 12 | -40 Å | -25 Å | -10 Å | -5 Å | 0 Å | 3.25 Å | 6.5 Å | 9.5 Å | 12.5 Å | 26.25 Å | 40 Å |

Only experimental data from single channel I-V relationships found in Heginbotham et al. (43 mM, 73 mM, 206 mM, 325 mM, 605 mM, 1.15 M)[14] and microscopic reversibility constraints were included in the loss function of the optimization procedure. Further details of the optimization procedure can be found in Weckel-Dahman et al.[8]

### S2. SUPPLEMENTAL RESULTS

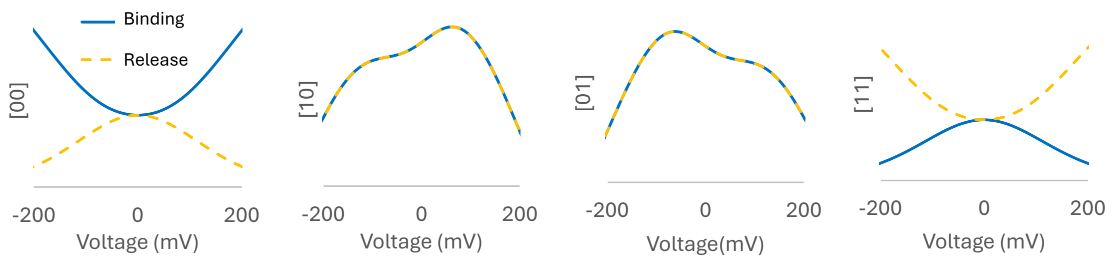


Figure S1. Population shifts of conductance shape 2 for populations shown in Figure 3C binding limited (blue) and Figure 3D release limited (yellow dashed). Initial population rises for state [10] and [01] become apparent when viewed in this limited voltage range. Population scale is not normalized.

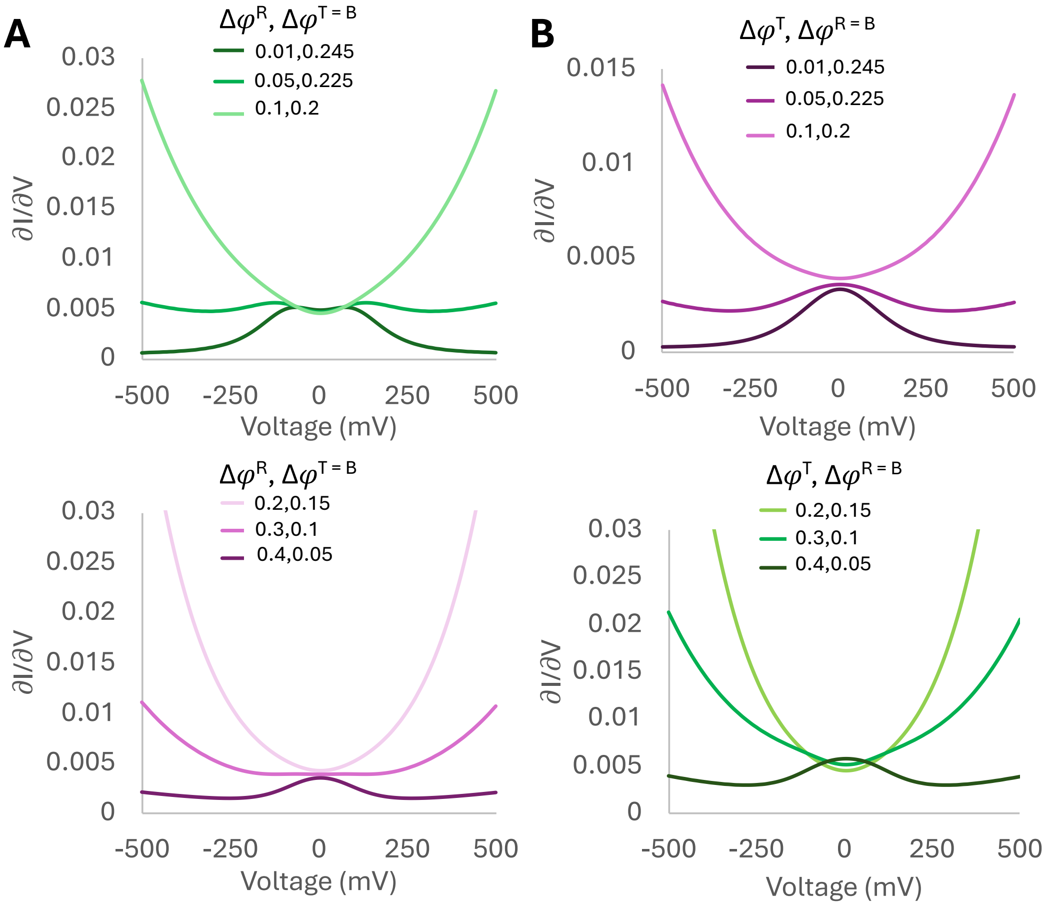


Figure S2. Scanning through the $\Delta\varphi$ of A) release and B) transfer. Curves equivalent to A were found when scanning the $\Delta\varphi$ of binding. The FLS gradually shifts from being binding and/or release limited (dark green → light green) to transfer limited (light purple → dark purple). Darker colors represent a system that is more limited.

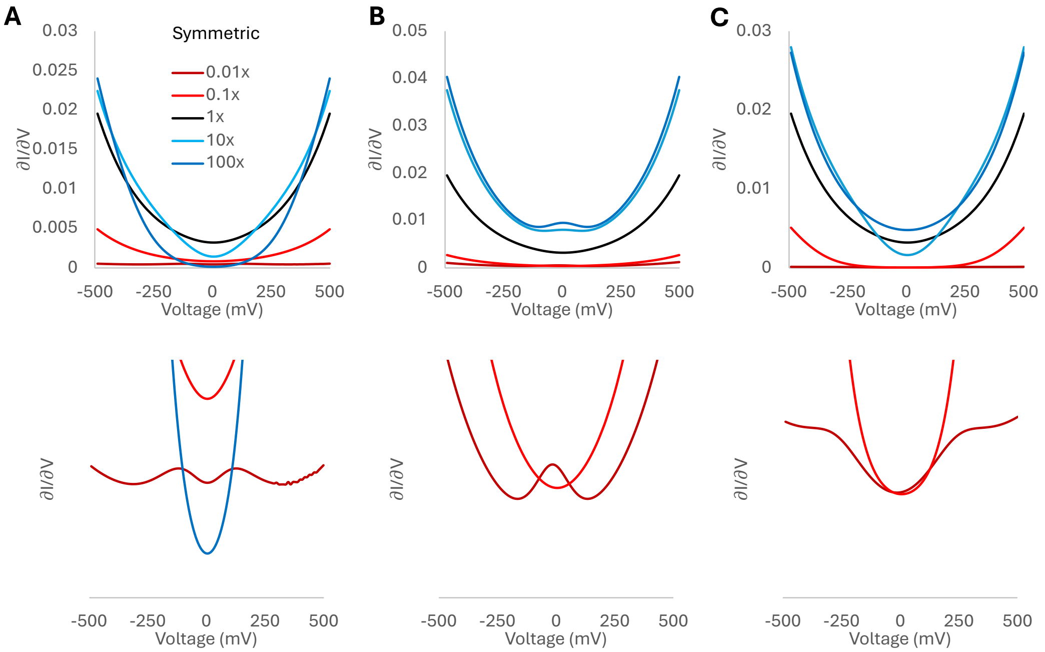


Figure S3. Effects of relative increases or decreases in the A) binding, B) transfer, and C) release rate constants on the conductance of a 3-site perfectly symmetric model.

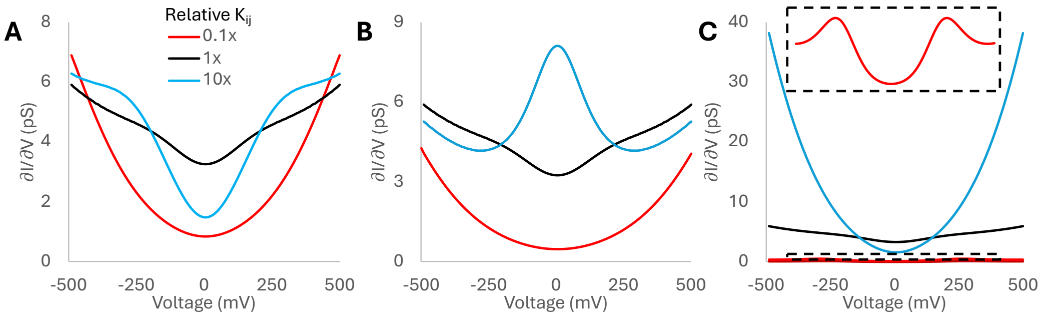


Figure S4. Effects of relative increases or decreases in the A) binding, B) transfer, and C) release rate constants on the conductance of a 3-site mirror-symmetric model with a release-limited VLS ($\Delta\varphi$ of $\pm$ 0.05).

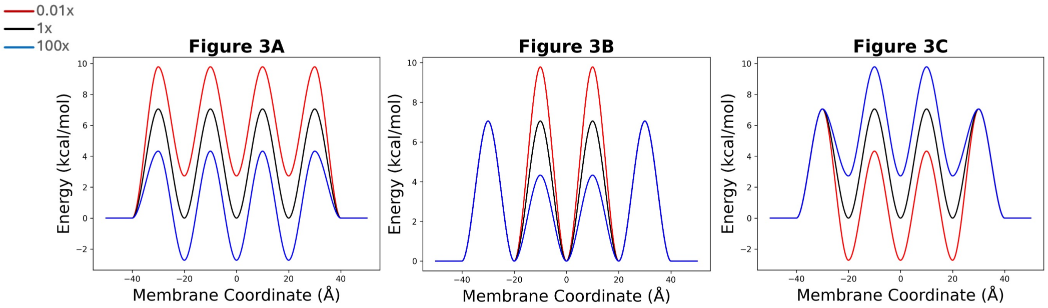


Figure S5. PMF curves for the systems shown in main text Figure 3.

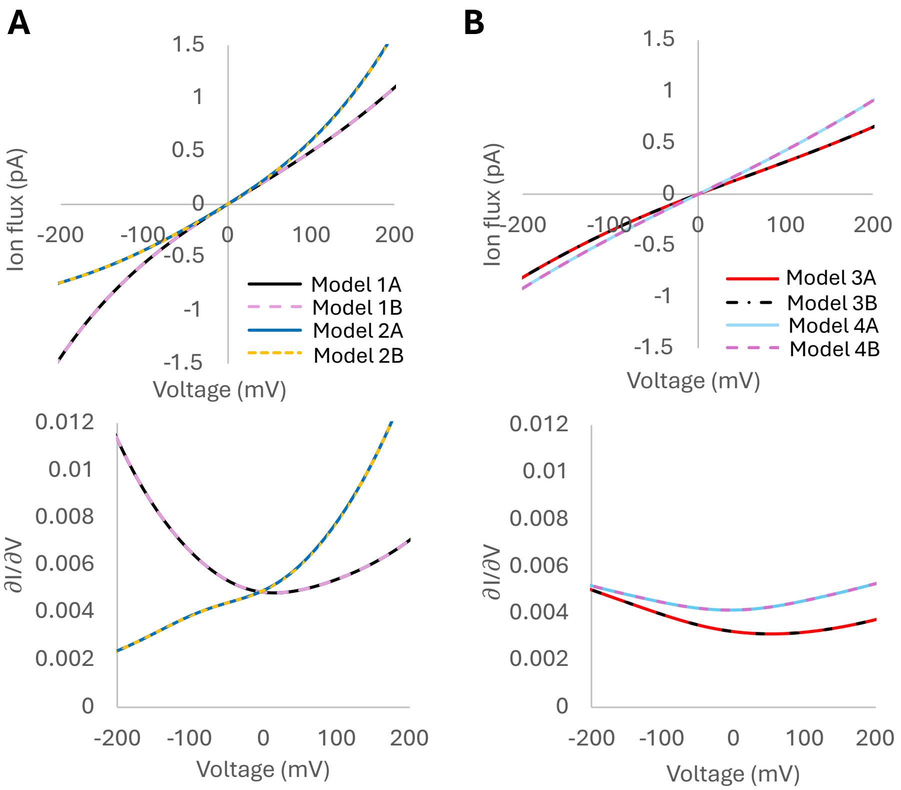


Figure S6. Ion flux (top) and conductance (bottom) for 2-site (A) and 3-site (B) models. Models are insensitive to $\Delta\varphi$ location as long as the conductance shape is maintained (see Tables S5 and S6).

Table S5. Voltage sensitivities for the 2-site models in Figure S3

| | $\|\Delta\varphi\|B_{out}$ 00 -> 10 | $\|\Delta\varphi\|R_{in}$ 10 -> 00 | $\|\Delta\varphi\|R_{out}$ 01 -> 00 | $\|\Delta\varphi\|B_{in}$ 00 -> 01 | $\|\Delta\varphi\|T_{in}$ 01-> 10 | $\|\Delta\varphi\|T_{out}$ 10 -> 01 |
|---|---|---|---|---|---|---|
| **Model 1A** | 0.1 | 0.2 | 0.2 | 0.2 | 0.15 | 0.15 |

| | | | | | | |
|---|---|---|---|---|---|---|
| **Model 1B** | 0.2 | 0.2 | 0.1 | 0.2 | 0.15 | 0.15 |
| **Model 2A** | 0.2 | 0.2 | 0.29 | 0.01 | 0.15 | 0.15 |
| **Model 2B** | 0.29 | 0.01 | 0.2 | 0.2 | 0.15 | 0.15 |

B/T/R stand for binding/transfer/release and in/out stand for inward and outward flux.

Table S6. Voltage sensitivities for the 3-site models in SI Figure S3

| | $\|\Delta\varphi\|T_{out}$ 100 -> 010 | $\|\Delta\varphi\|T_{in}$ 010 -> 100 | $\|\Delta\varphi\|T_{out}$ 010 -> 001 | $\|\Delta\varphi\|T_{in}$ 001 -> 010 | $\|\Delta\varphi\|$ B/R |
|---|---|---|---|---|---|
| **Model 3A** | 0.1 | 0.2 | 0.2 | 0.2 | 0.075 |
| **Model 3B** | 0.2 | 0.2 | 0.1 | 0.2 | 0.075 |
| **Model 4A** | 0.2 | 0.2 | 0.29 | 0.01 | 0.075 |
| **Model 4B** | 0.29 | 0.01 | 0.2 | 0.2 | 0.075 |

B/T/R stand for binding/transfer/release and in/out stand for inward and outward flux.

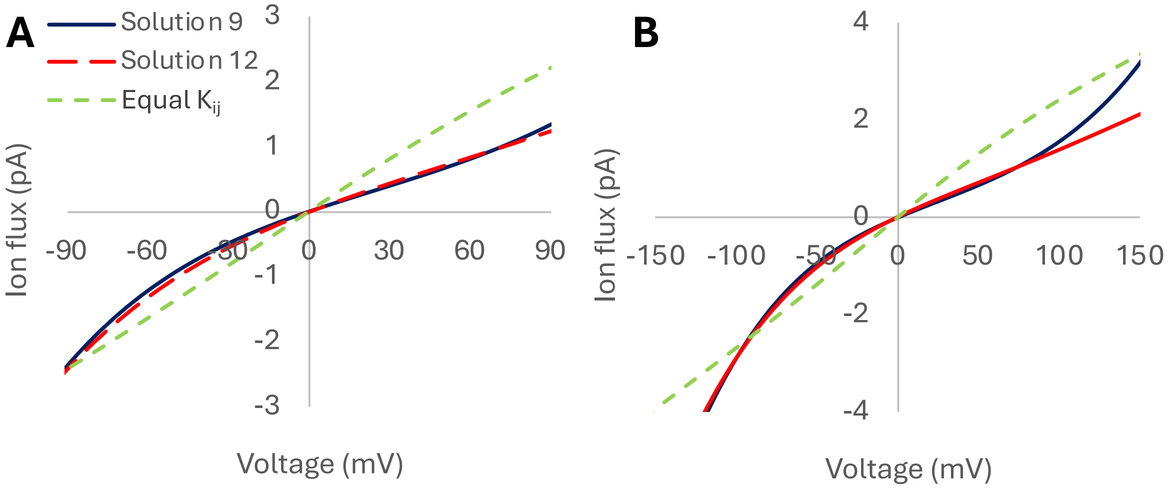


Figure S7. Solutions 9 (solid black) and 12 (red dashed) flux compared to the flux generated with symmetric rate constants (green dashed). Since the two solutions have identical voltage sensitivities, the differences in rectification are a result of differences in rate constants. Note the overlap in the voltage range included in optimization (A) that is quickly lost when extending to a larger voltage range (B).
